# Genome location dictates the transcriptional response to PolC-inhibition in *Clostridium difficile*

**DOI:** 10.1101/362137

**Authors:** Erika van Eijk, Ilse M. Boekhoud, Ed J. Kuijper, Ingrid M.J.G. Bos-Sanders, George Wright, Wiep Klaas Smits

## Abstract

*Clostridium difficile* is a potentially lethal gut pathogen that causes nosocomial and community acquired infections. Limited treatment options and reports of reduced susceptibility to current treatment emphasize the necessity for novel antimicrobials. The DNA-polymerase of gram-positive organisms is an attractive target for the development of antimicrobials. ACX-362E (*N*^2^-(3<,4-Dichlorobenzyl)-7-(2-[1-morpholinyl]ethyl)guanine; MorE-DCBG) is a DNA polymerase inhibitor in pre-clinical development as a novel therapeutic against *C. difficile* infection. This synthetic purine shows preferential activity against *C. difficile* PolC over those of other organisms *in vitro* and is effective in an animal model of *C. difficile* infection. In this study we have determined its efficacy against a large collection of clinical isolates. At concentrations below the minimal inhibitory concentration, the presumed slowing (or stalling) of replication forks due to ACX-362E leads to a growth defect. We have determined the transcriptional response of *C. difficile* to replication inhibition and observed an overrepresentation of up-regulated genes near the origin of replication in the presence of PolC-inhibitors, but not when cells were subjected to sub-inhibitory concentrations of other antibiotics. This phenomenon can be explained by a gene dosage shift, as we observed a concomitant increase in the ratio between origin-proximal versus terminus-proximal gene copy number upon exposure to PolC-inhibitors. Moreover, we show that certain genes differentially regulated under PolC-inhibition are controlled by the origin-proximal general stress response regulator sigma factor B. Together, these data suggest that genome location both directly and indirectly determines the transcriptional response to replication inhibition in *C. difficile*.

## Background

*Clostridium difficile* (*Clostridioides difficile* (1)) is a gram-positive, anaerobic bacterium that can asymptomatically colonize the intestine of humans and other mammals (2-4). However, when the normal flora is disturbed *C. difficile* can overgrow and cause fatal disease, as has been dramatically demonstrated in the Stoke Mandeville Hospital outbreaks in 2004 and 2005 (5). The ability to form highly resistant endospores coupled to its extensive antibiotic resistance have contributed to its success as a nosocomial and community acquired pathogen (2-4). Recent years have seen an increase in the incidence and severity of *C. difficile* infections (CDI), due to the emergence of certain PCR ribotypes (3, 6). Antibiotic use is a well-established risk factor for CDI (7), and the emergence of the epidemic PCR ribotype 027 has been linked to fluoroquinolone resistance (8). At present two antibiotics, metronidazole and vancomycin are commonly used to treat CDI and a third, fidaxomicin, is indicated for the treatment of relapsing CDI (9, 10). Clearly, limited treatment options and reports of reduced susceptibility to current treatment (11-13) emphasize the necessity for the development of novel antimicrobials and a better understanding of tolerance and resistance towards existing therapeutics.

It is increasingly realized that off-target effects that occur when cells are exposed to antimicrobials can contribute to their efficacy, but also facilitate the emergence of tolerance and/or resistance (14). Antimicrobials may act as signaling molecules which modulate gene expression (14). Additionally, in particular those targeting DNA replication (such as polymerase inhibitors) can cause transcriptional effects as a result of differences in gene dosage (15).

The polymerase of gram-positive organisms is an attractive target for the development of novel antimicrobials (16). First, these PolC-type polymerases are absent from gram-negative organisms and humans (17, 18). HPUra, one of the first such compounds, is therefore highly active against a wide range of gram-positive bacteria, but does not affect gram-negative bacteria (17, 18). Template-directed elongation is blocked by the inhibitor through simultaneous binding to the cytosine of the DNA strand and near the active site of PolC. Second, compounds can be derived that have an increased specificity towards specific microorganisms. ACX-362E **(Figure 1)** is a compound in pre-clinical development as a novel therapeutic against *C. difficile* as it is shows preferential activity against *C. difficile* PolC over those of other organisms *in vitro* (19, 20), and will progress to clinical trials in the near future (Acurx Pharmaceuticals, personal communication). PolC-inhibitors can cause a stress response and cell death after prolonged exposure. In *Bacillus subtilis*, this stress is characterized by a combination of DNA damage (SOS) response, and an SOS-independent pathway dependent on the DNA replication initiator DnaA (21, 22). In *Streptococcus pneumoniae* cells, devoid of an SOS-response, competence for genetic transformation is induced upon replication stress (23). The response of *C. difficile* to this particular class of compounds is unknown.

**Figure 1.**
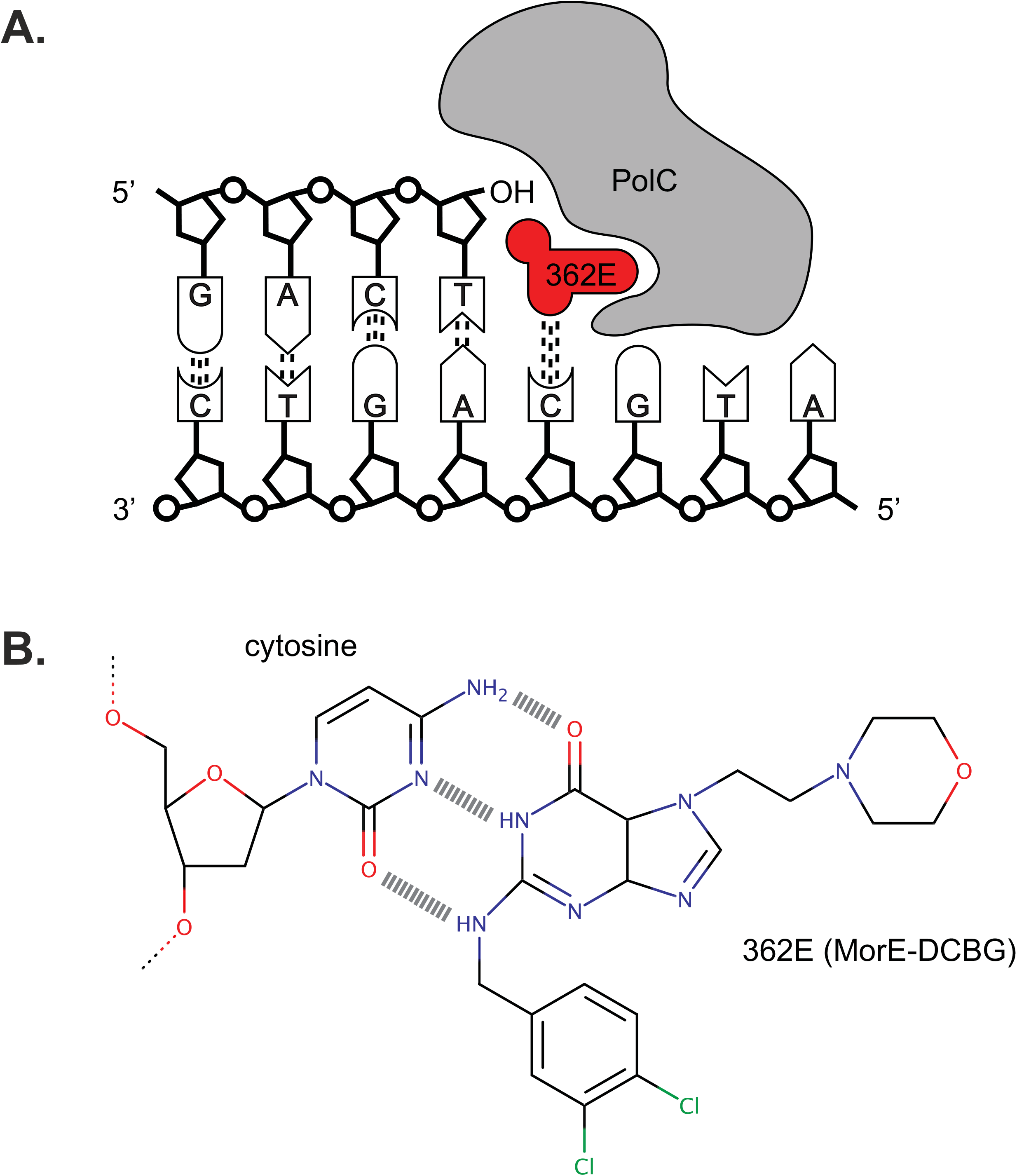
Mechanism of action of the PolC-inhibitors ACX-362E. **A.** Ternary complex of inhibitor ACX-362E, DNA, and PolC. **B.** H-bonding between inhibitor molecule ACX-362E and a cytosine residue of DNA.

In this study, we characterized aspects of the action of PolC-inhibitors towards *C. difficile*. Minimal inhibitory concentrations for HPUra and ACX-362E were determined using agar dilution for a large collection of clinical isolates. Next, we investigated the effects of sub-inhibitory levels of PolC-inhibitors on growth of *C. difficile* in liquid medium and performed RNAseq analyses to determine the transcriptional response to PolC-inhibitors in our laboratory strain 630Δ*erm*. Finally, marker frequency analysis and transcriptional reporters were used to provide a mechanistic explanation for the observed up-regulation of origin-proximal genes under conditions of replication inhibition.

## Results

### ACX-362E is a potent inhibitor of diverse clinical isolates of C. difficile

To date, reports on activity of PolC-inhibitors towards *C. difficile* are limited. For only 4 and 23 *C. difficile* strains minimal inhibitory concentrations were published (19, 20), and no analysis was performed on possible differences in efficacy between various phylogenetic groups (24, 25). Therefore, we assessed the sensitivity of a diverse collection of *C. difficile* clinical isolates towards PolC-inhibitors and determined if ACX-362E was indeed superior to the general PolC–inhibitor HPUra.

HPUra and ACX-362E were tested by the agar dilution method, according Clinical and Laboratory Standards Institute (CLSI) guidelines for testing of antimicrobial susceptibility of anaerobes (26, 27), against 364 *C. difficile* clinical isolates collected earlier in the framework of a pan-European study (6, 28).

We found that ACX-362E (MIC_50_: 2 µg/ml; MIC_90_: 4 µg/ml) demonstrates lower inhibitory concentrations than the general gram-positive PolC-inhibitors HPUra (MIC_50_: 16 µg/ml; MIC_90_: 32 µg/ml) (**Figure 2, Supplemental Table 1**), consistent with previous *in vitro* activities against purified PolC (19). We observed no significant difference in ACX-362E susceptibility between clades **(Table 1)** and the different PCR ribotypes demonstrated a similar distribution in MIC values (data not shown). No growth at the highest concentration of compounds tested (resistance) for either one of the PolC-inhibitors was observedamong the clinical isolates tested (N=363). Notably, we observed only a 2-fold difference in MIC_50_ and MIC_90_, indicating that the compounds have similar activity against nearly all strains. In contrast, the gram-negative obligate anaerobe *Bacteroides fragilis* was resistant to both polymerase inhibitors under the conditions tested (MIC >265 µg/ml), as expected for an organism lacking PolC. The gram-positive bacterium *Staphylococcus aureus*, which was included as a control for the activity of HPUra against this group of bacteria (16, 29), was sensitive to both HPUra and ACX-362E, with MIC values of 2 μg/mL and 1 μg/mL, respectively.

**Figure 2.**
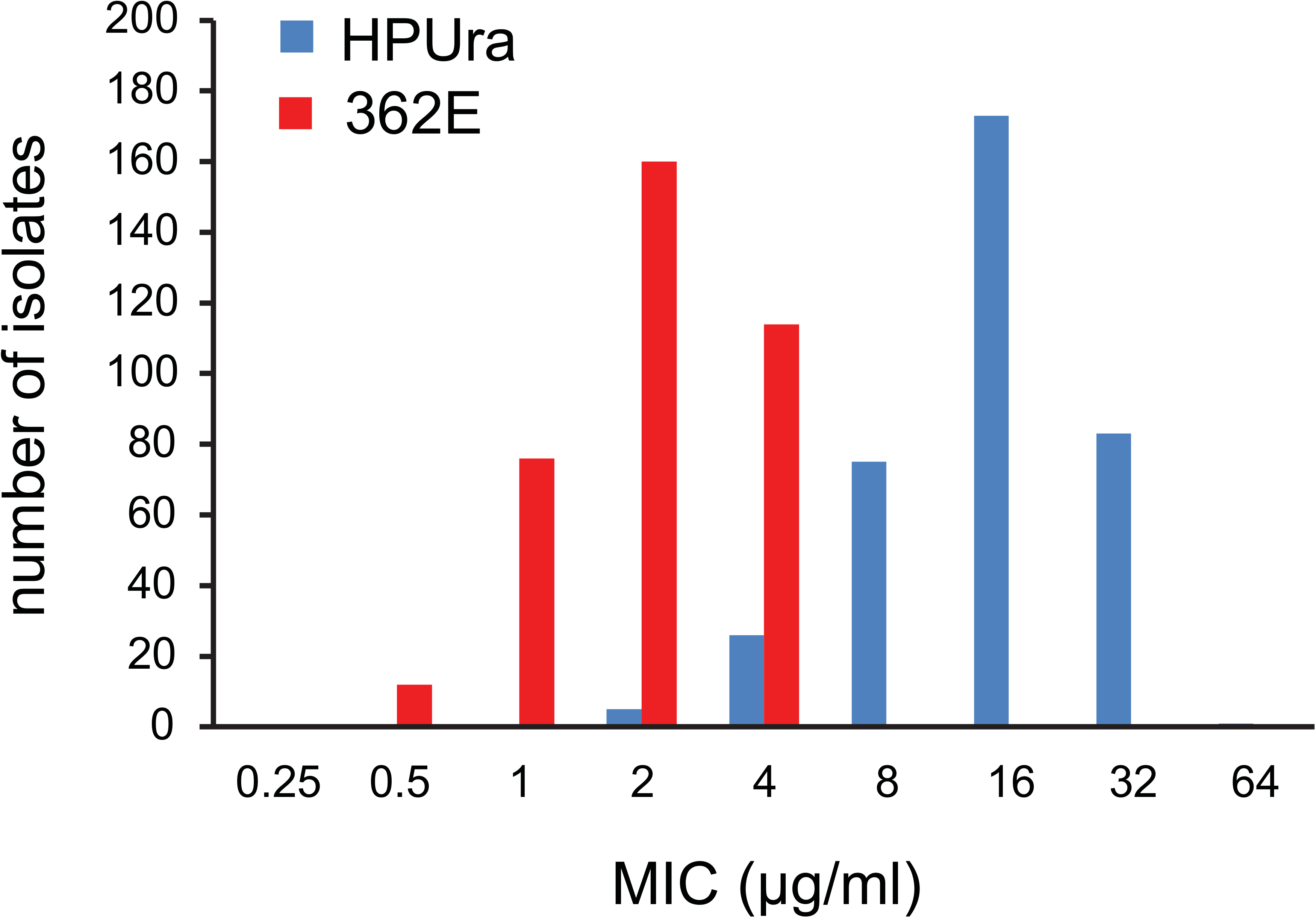
Minimal inhibitory concentrations of PolC-inhibitors. MIC was determined by agar dilution according to CLSI standards (26) and is expressed in μg/mL. The distribution in the MIC for the collection of clinical isolates (N=363) is given for the PolC-inhibitors HPUra (blue) and ACX-362E (red).

**Table 1.** Minimal inhibitory concentrations of PolC-inhibitors towards *C. difficile* stratified by clade.

We conclude that ACX-362E is highly active against diverse clinical isolates of *C. difficile* and resistance is not a concern in currently circulating strains.

### Treatment with ACX-362E leads to a pleiotropic transcriptional response

In order to determine the transcriptional response of PolC-inhibitors, we established the optimal concentration of both inhibitors which affected growth of *C. difficile* in liquid medium. The laboratory strain *C. difficile* 630*Δerm* (PCR ribotype 012, MLST Clade 1)(30, 31) was grown in medium with varying amounts of HPUra (10-40 μg/mL) or ACX-362E (0.25-8 μg/mL). We note that concentrations up to the MIC_90_ (as determined by agar dilution) did not lead to a complete growth arrest in liquid medium in the time course of the experiment (**Supplemental Figure 1**). A difference between MIC values from agar dilution and (micro)broth has been observed before (32). The growth kinetics of *C. difficile* under the influence of varying concentrations of HPUra was marginally affected when using concentrations from 10-40 µg/ml, at >80 percent of the non-treated culture. Growth kinetics of cultures containing PolC-inhibitor ACX-362E 1-8 µg/ml were similar and resulted in -30-40 percent reduced growth compared to the untreated culture. For subsequent experiments we used concentrations of PolC-inhibitors that were at or close to the MIC_90_ (**Table 1**) and that result in a maximum reduction in growth of 30% compared to a non-treated culture (**Supplemental Figure 1**).

Above, we established that growth of *C. difficile* is partially inhibited at certain concentrations of PolC-inhibitors. Slowing down or stalling of replication forks might lead to a stressed state, as observed for other organisms (22, 23). As nothing is known about the effect of replication inhibition on the physiology of *C. difficile*, we determined the transcriptional response to replication inhibition by sub-MIC levels of PolC-inhibitors through strand-specific RNA sequencing (RNA-Seq).

*C. difficile* 630*Δerm* was grown for 5h in medium with HPUra (35 μg/mL) or ACX-362E (4 µg/mL) starting from an OD_600_ of 0.05 after which cells were harvested for RNA isolation. Purified RNA was converted to cDNA and used for RNA-Seq as described in the Materials and Methods. For ACX-362E, 722 genes were differentially expressed, of which 438 genes were up-regulated and 284 genes were down-regulated. The number of differentially expressed genes in HPUra treated samples was approximately 2-fold lower: 360, of which 124 genes were up-regulated and 236 genes were down-regulated. The full list of differentially regulated genes is available as **Supplementary Table 2** and the top 10 of up- and down-regulated genes are shown in **Table 2** (HPura) and **Table 3** (ACX-362E). Here, we highlight three aspects of the results.

**Table 2.** List of the genes most highly up- and down-regulated in the presence of HPUra compared to non-treated cells.

**Table 3.** List of the genes most highly up- and down-regulated in the presence of ACX-362E compared to non-treated cells.

First, we performed a Gene Set Enrichment Analysis (GSEA)(33) via the Genome2D web server (34) using the locus tags of the differentially regulated genes (**Supplementary Table 1**) as input. Among the genes up-regulated by ACX-362E, there is a strong overrepresentation of those involved in translation, ribosome structure and ribosome biogenesis. Not unexpectedly, also replication, recombination and repair processes are affected. This suggests that genes from these pathways show a coordinated response in the presence of ACX-362E. Among the genes down-regulated in the presence of ACX-362E, the levels of significance for specific processes are generally much lower, suggesting that there is a more heterogeneous response among genes from the same pathway. Nevertheless, metabolic pathways (especially carbon metabolism and coenzyme A transfer) and tellurite resistance were found to be significantly affected. Strikingly, a GSEA analysis on lists of genes that are differentially expressed in the presence of HPUra revealed similar processes to be affected.

The findings from the GSEA analysis prompted us to evaluate the overlap in the lists of differentially regulated genes between the ACX-362E and HPUra datasets in more detail. If both compounds act via a similar mechanism, we expect a conserved response. Indeed, we observe that >90% of the genes that are up-regulated in the presence of HPUra compared to the non-treated condition, are also identified as up-regulated in the presence of ACX-362E (**Figure 3A**). Though the overlap is not as strong for the down-regulated genes, we find that >30% of the genes affected by HPUra are also identified as affected by ACX-362E (**Figure 3B**). Notably, the directionality of the response is conserved, as no genes were found to be up-regulated in one but down-regulated in the other condition. Based on these observations, we believe that the differentially expressed genes identified in this study are representative for a typical response to inhibition of PolC in *C. difficile*.

**Figure 3.**
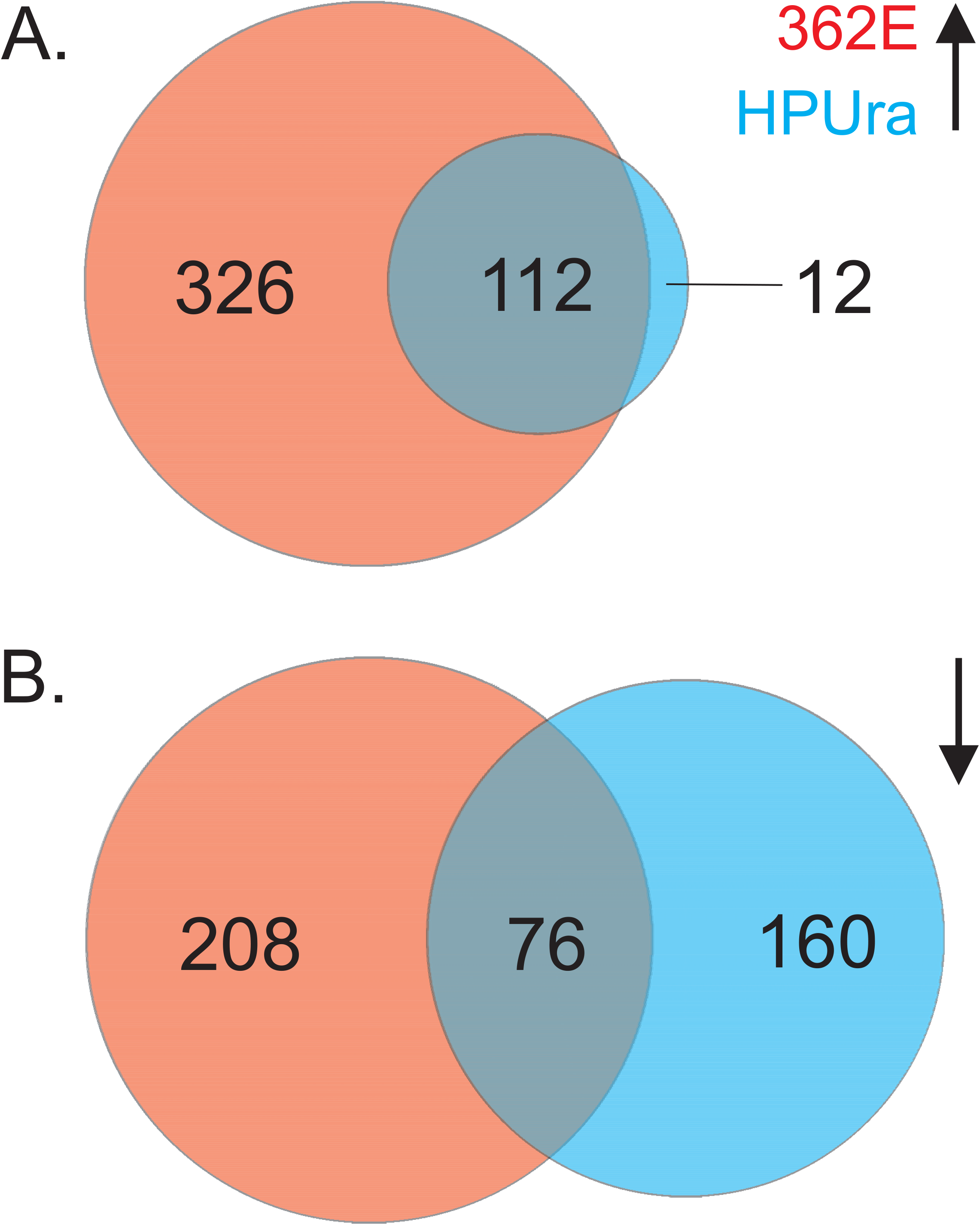
Overlap in the transcriptional response to different PolC-inhibitors. **A.** Venn diagram of the number of genes up-regulated in the presence of ACX-362E (red), in the presence of HPUra (blue) or under both conditions (overlapping region). **B.** Venn diagram of the number of genes down-regulated in the presence of ACX-362E (red), in the presence of HPUra (blue) or under both conditions (overlapping region). The size of the circles is proportional to the number of genes that showed differential expression.

Finally, we related the changes in transcription to genome location. *C. difficile* has a single circular chromosome and one origin of replication (*oriC*) from which the process of DNA replication occurs bi-directionally towards the terminus (*terC*) (**Figure 4A**). Though neither *oriC* nor *terC* has been definitively defined for *C. difficile*, it is assumed that *oriC* is located at or near *dnaA* (CD0001; CD630DERM_RS00005). The terminus region is generally located at the inflection point of a GC skew ([G - C]/[G + C]) plot. Such a plot places the *terC* region around 2.2Mb from CD0001, near the CD1931 (CD630DERM_RS10465) open reading frame (**Figure 4A**) (35). We noted that the differential expression appeared to correlate with genome location (**Table 2, Table 3** and **Supplemental Table 2**): many of the up-regulated genes have either low or high gene identifiers (CD numbers) indicative of an origin proximal location and, conversely, many of the down-regulated genes appear to be located away from *oriC*. Though this correlation is not absolute, we observed a clear trend when plotting the mean fold-change against genome location for all genes (**Figure 4B**).

**Figure 4.**
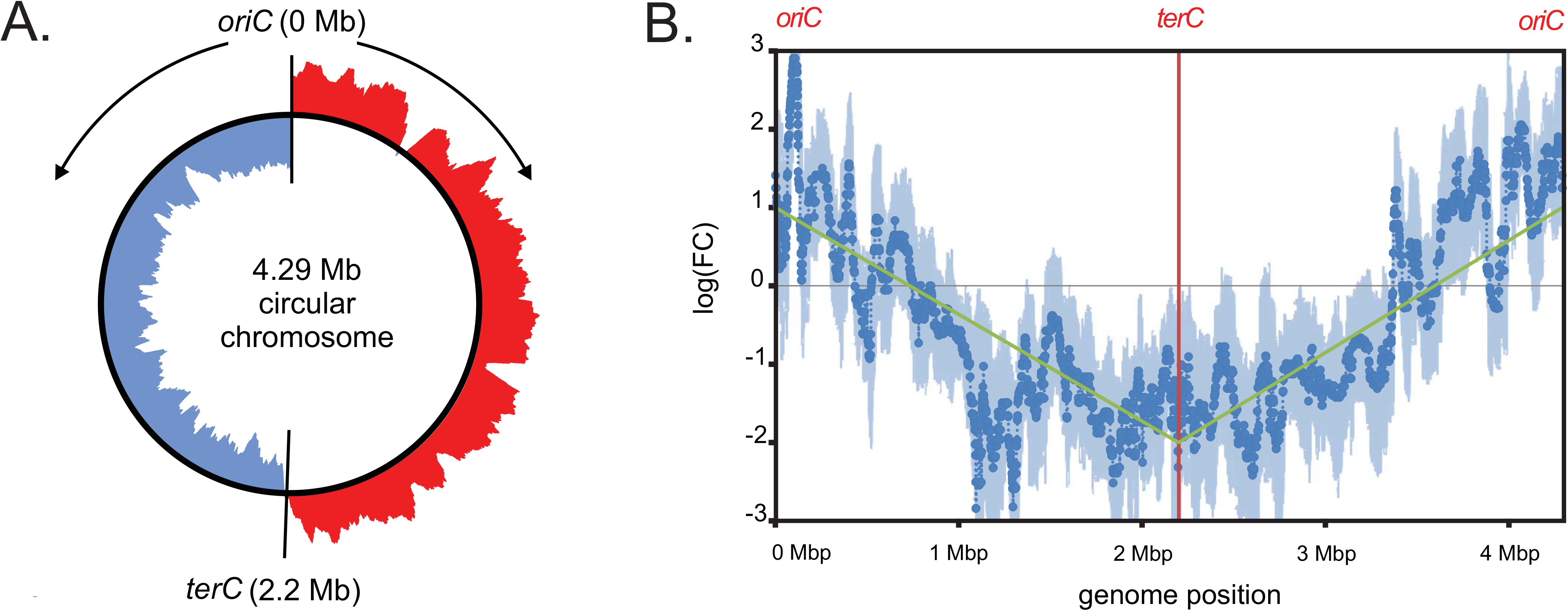
Genome location correlates with differential expression upon PolC inhibition. **A.** Schematic representation of the chromosome of *C. difficile*. Higher than average GC skew ([G - C]/[G + C]) (red) and lower than average GC skew (blue) were calculated with DNAplotter (72). Vertical lines indicate the position of the predicted origin (*oriC*) and terminus (*terC*) of replication. Arrows indicate the direction of replication. **B.** Sliding window analysis (bins of 51 loci, stepsize 1) of the median log fold change (FC) projected on a linear genome map. The *oriC* of the circular chromosome is located on either size of the linear graph (0/4.29Mb), whereas *terC* is indicated with a vertical red line. The trend in log(FC) is highlighted using a green line. Light blue shading indicates the median absolute deviation of the mean (23).

Overall, our data shows that inhibition of DNA replication by PolC-inhibitors causes a consistent and pleiotropic transcriptional response that is at least in part is directly dictated by genome location.

### Gene dosage shift occurs at sub-inhibitory concentration of ACX-362E PolC-inhibitor

A possible explanation for the relative up-regulation of *oriC*-proximal genes and down-regulation of *terC*-proximal genes is a gene dosage shift (36-38), due to the fact that PolC-inhibition slows down replication elongation but does not prevent re-initiation of DNA replication (23, 39). To determine if this in fact occurs in *C. difficile* when replication elongation is inhibited, we performed a marker frequency analysis (MFA) to determine the relative abundance of an origin proximal gene relative to terminus proximal gene on chromosomal DNA isolated from treated and untreated cells.

We designed qPCR probes against the CD0001 and CD1931 regions, representing *oriC* and *terC*, respectively (31, 35). Using these probes, we could show that *C. difficile* demonstrates multi-fork replication in exponential growth phase and that the MFA assay detects the expected decrease in *oriC*:*terC* ratio when cells enter stationary growth phase (data not shown). Next, we analyzed the effects of PolC-inhibitors on the *oriC*:*terC* ratio. When cells were treated with HPUra (35 μg/mL), the MFA showed a modest increase in *oriC:terC* ratio of 2,6-fold compared to untreated cells. However, when cells were treated with ACX-362E (4 µg/ml), the MFA showed a >8-fold increase in the *oriC*:*terC* ratio compared to untreated cells. In contrast, such an increase was not observed for cells treated with metronidazole (0.25 ug/mL; a DNA damaging agent), fidaxomicin (0.00125 ug/mL; an RNA polymerase inhibitor) or surotomycin (0.625 ug/mL; a cell-wall synthesis inhibitor) (**Figure 5**) or chloramphenicol (2 μg/mL; a protein synthesis inhibitor)(**Supplemental Figure 2**).

**Figure 5.**
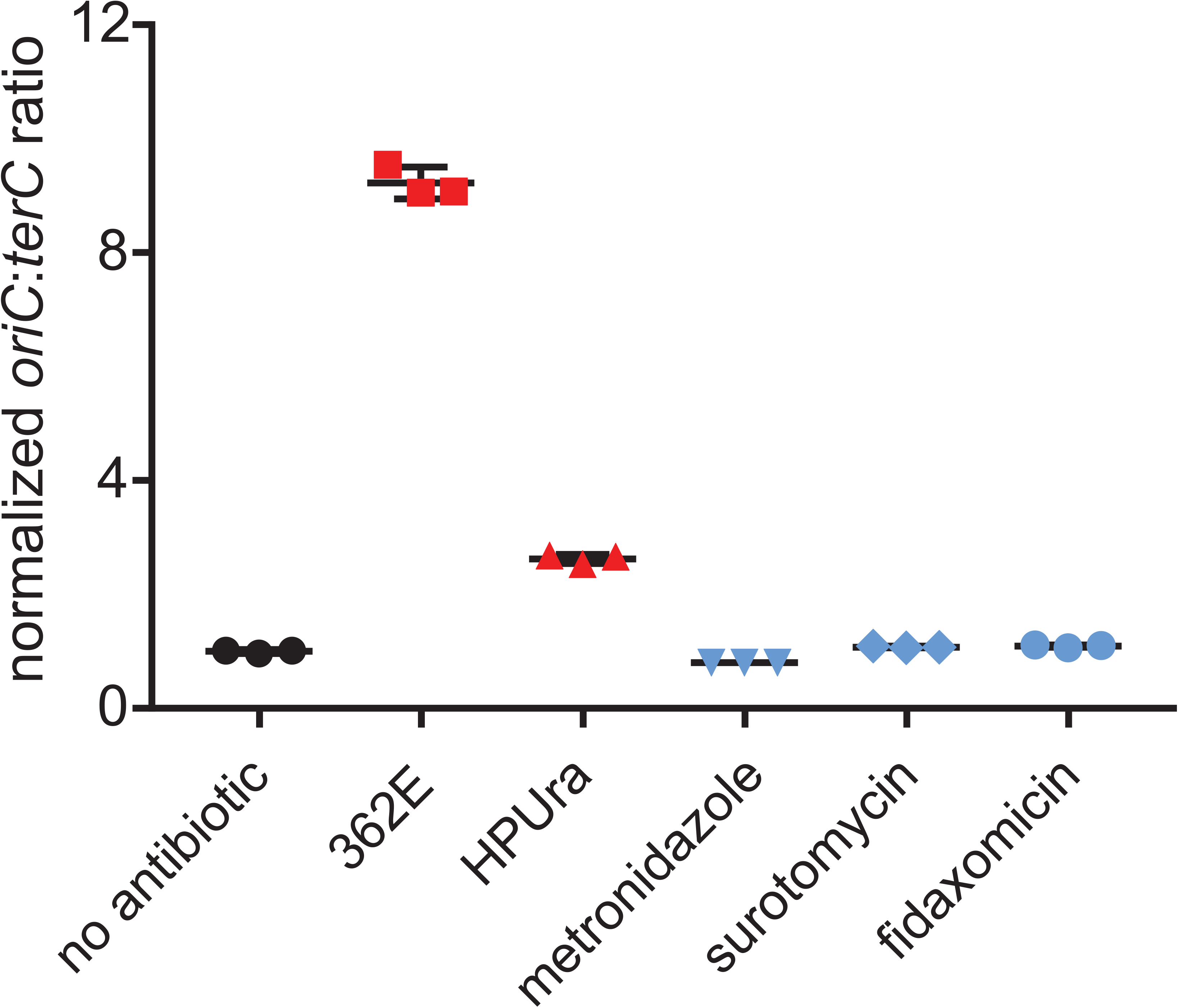
Polymerase inhibitors lead to an increase in *oriC:terC* ratio. A marker frequency analysis of the effects of sub-inhibitory amounts of polymerase inhibitors (red; HPUra: 35 μg/mL; ACX-362E: 4 μg/mL) and three antibiotics with different modes of action (blue; metronidazole: 0.25 μg/mL; fidaxomicin: 0.00125 ug/mL, surotomycin: 0.625 μg/mL) compared to non-treated cells (black). Datapoints are averages of technical replicates (N=3). Black lines behind the datapoints indicate the average of the biological replicates (N=3) and whiskers indicate the standard deviation of the mean. Data have been normalized compared to the non-treated control. The mean of HPUra and ACX-362E treated samples is statistically different from the other samples (p<0.0001).

We conclude that inhibition of PolC-activity, but not the actions of any of the other tested antimicrobials, leads to a gene dosage shift in *C. difficile*.

### *Origin proximal* sigB *contributes to the transcriptional response*

Elegant work in *S. pneumoniae* has shown that the transcriptional response to replication inhibition can also be affected by origin-proximal regulators that respond to the gene dosage effect (23). In *C. difficile* the gene encoding the general stress response sigma factor σ^B^ (*sigB*, CD0011) is located close to the origin of replication (40). We wondered whether this regulator contributes to the transcriptional effects observed in our studies.

First, we compared the list of differentially regulated genes from our study (**Supplemental Table 2**) to those under the control of σ^B^ (41). In contrast to most anaerobic gram-positive organisms, *C. difficile* encodes a homolog of the general stress response sigma factor, σ^B^ (40, 42). A transcriptome analysis comparing *sigB* mutant versus wild type cells was recently published (41). Strikingly, we find ~40% of the genes (21/58) identified as involved in stress response under the control of σ^B^ to be differentially expressed in our ACX-362E dataset (**Supplemental Table 3**). Similarly, we observed that 7/20 (~35%) of the genes containing a transcriptional start site with a σ^B^ consensus sequence are differentially expressed in our ACX-362E dataset (**Supplemental Table 3**). These data suggest that the response to DNA replication inhibition is at least partially dependent on σ^B^.

To demonstrate that exposure to ACX-362E causes a transient up-regulation of *sigB*, we constructed a reporter fusion of the secreted luciferase reporter sLuc^opt^ with the predicted promoter of the *sigB* operon (P*_cd0007_*). We monitored luciferase activity of a strain harboring a plasmid containing this reporter fusion (WKS2003) after dilution of an overnight culture into fresh medium with or without ACX-362E (**Figure 6**). In non-treated cells, expression from the *sigB* promoter is relatively stable over the course of 5.5h. By contrast, luciferase activity strongly increases from 1.25h to 3h after inoculation into medium with ACX-362E. These data show that exposure to ACX-362E transiently induces transcription of the *sigB* operon.

**Figure 6.**
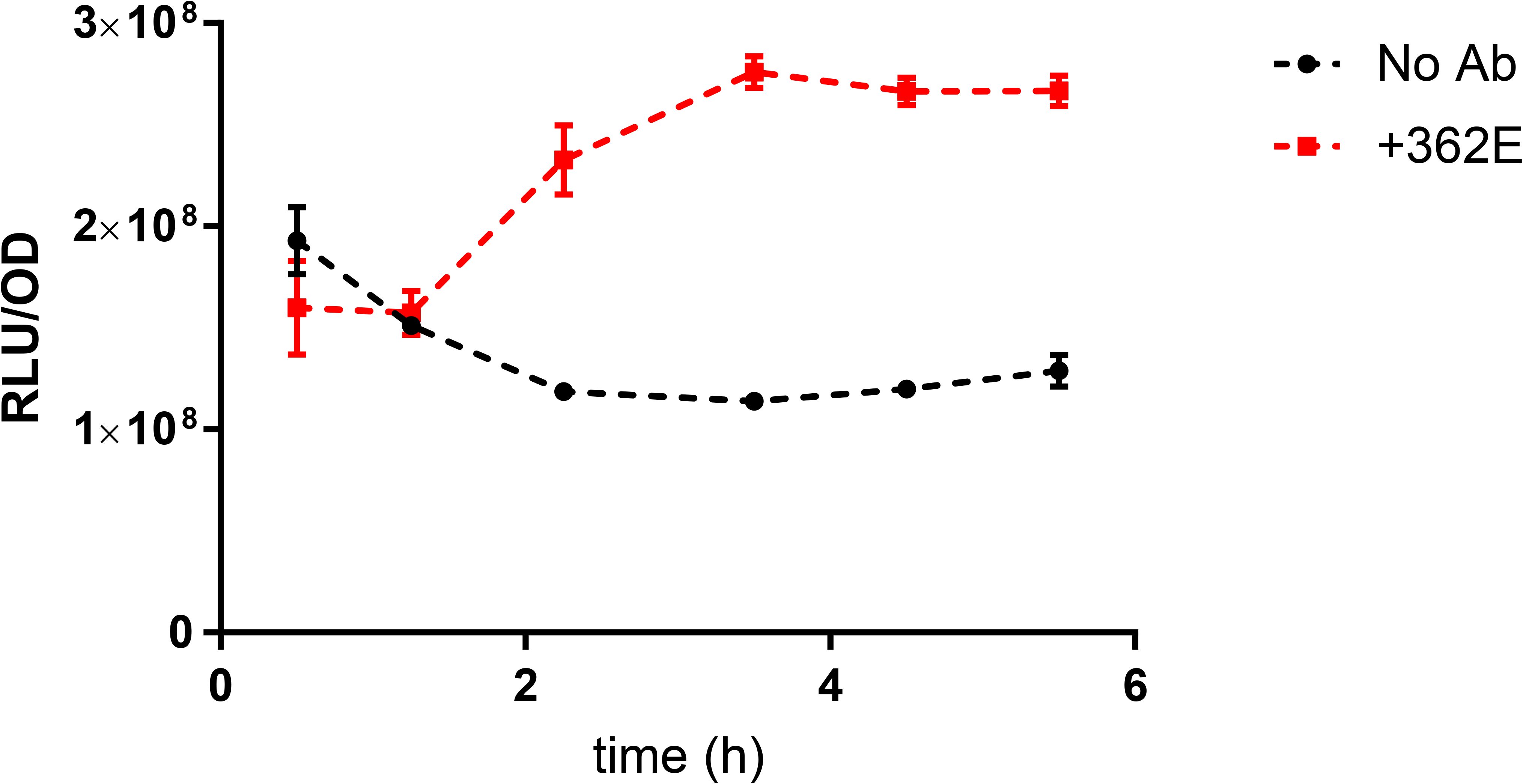
The *sigB* operon is transiently induced upon exposure to ACX-362E. The putative promoter of the *sigB* operon (41) was fused transcriptionally to a plasmid-based luciferase reporter (44). Luciferase activity was measured regularly between 30 mins and 5.5h of growth in liquid medium in the presence (+362E) or absence (no Ab) of polymerase inhibitor ACX-362E.

Next, a *sigB* mutant was constructed using Allele-Coupled Exchange (43) as described in the Materials and Methods. The chromosomal deletion of *sigB* and absence of σ^B^ protein was verified by PCR and Western Blot, respectively (**Supplemental Figure 3**). To directly demonstrate a role for *sigB* in the regulation of genes with altered transcription upon PolC-inhibition, we fused the predicted promoter regions of selected genes to a secreted luciferase reporter (44) and evaluated luminescence in wild type and *sigB* mutant backgrounds, after 5-hour growth in the presence and absence of 4 µg/mL ACX-362E. All genes tested demonstrated a significant increase in promoter activity in a wild type background in the presence of ACX-362E compared to the non-treated control, validating the results from the RNA-Seq analysis (**Figure 7**). Three distinct patterns were observed. For CD0350 (encoding a hypothetical protein) and CD3963 (encoding a putative peptidoglycan-binding exported protein) there was virtually no expression in a *sigB* mutant background (**Figure 7A** and **B**). We conclude that these genes are strictly dependent on σ^B^ for their expression under the conditions tested. CD3614 (encoding a hypothetical protein) shows a basal level of expression, but no significant increase in transcription levels in the presence of ACX-362E in a *sigB* mutant background compared to the non-treated control (**Figure 6C**). This suggests that the transcriptional up-regulation under these conditions is σ^B^ dependent, and indicates that the basal level of transcription observed is likely independent of σ^B^. Finally, CD3412 (*uvrB*, encoding a subunit of an excinuclease) shows reduced transcriptional up-regulation in ACX-362E treated cells compared to non-treated controls (**Figure 6D**). Thus, the transcription of this particular gene under conditions of ACX-362E exposure is brought about by both a σ^B^-dependent and a σ^B^-independent regulatory pathway.

**Figure 7.**
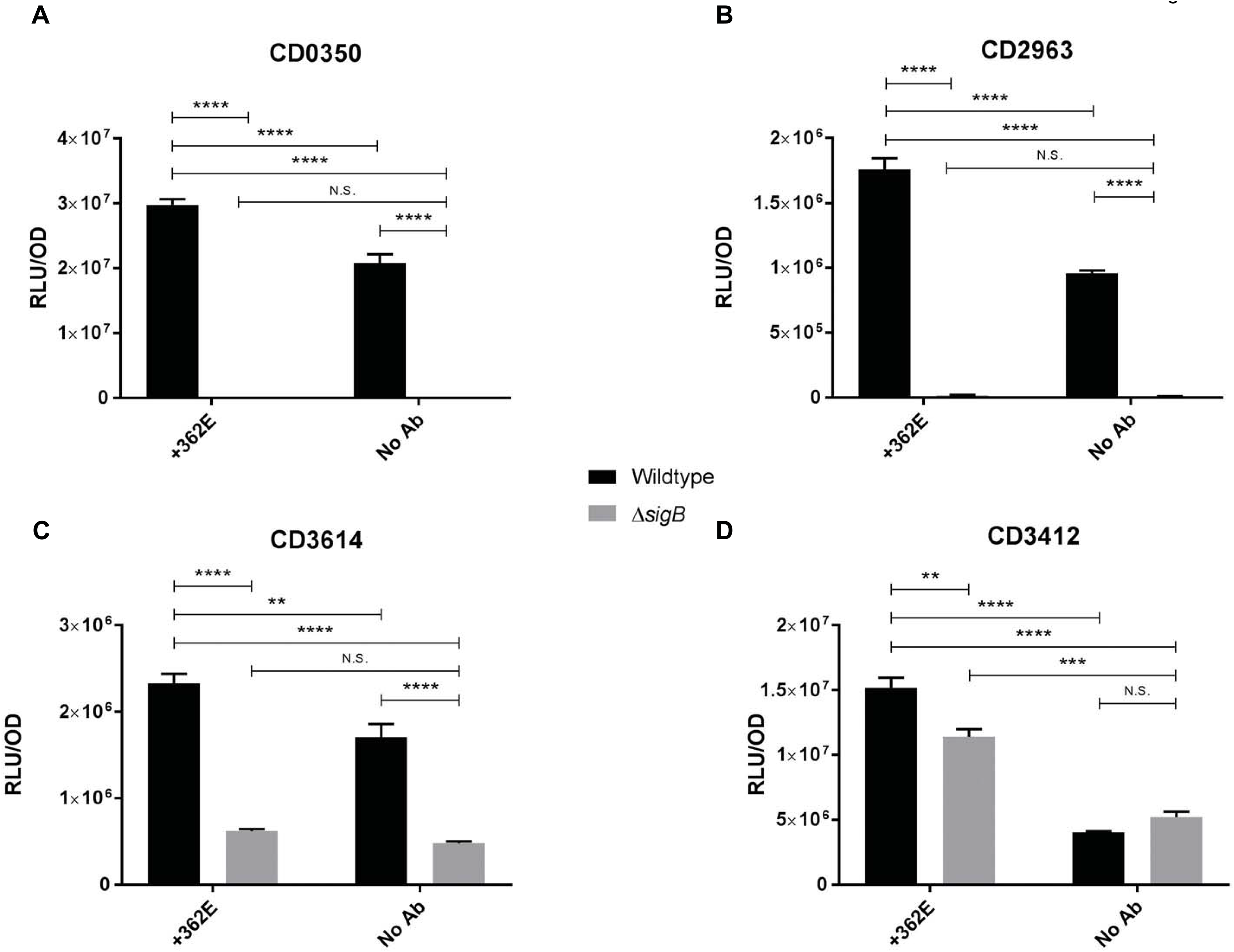
Genes differentially expressed due to polymerase inhibitors are regulated by σ^B^. The putative promoters of the indicated genes were fused transcriptionally to a plasmid-based luciferase reporter
(44). Luciferase activity was measured after 5 h of growth in liquid medium in the presence (+362E) or absence (no Ab) of polymerase inhibitor ACX-362E. N.S. = non-significant, *=p<0.05, **=p<0.005, ***=p<0.0005, ****=p<0.00005. **A**. P_CD350_-*sLuc*^opt^. **B**. P_CD2963_-*sLuc*^opt^ **C**. P_CD3614_-*sLuc*^opt^ **D**. P_CD3412_-*sLuc*^opt^.

Together, these results demonstrate that *sigB* controls the expression of at least a subset of genes that are up-regulated under PolC-inhibition

## Discussion

### Resistance to PolC-inhibitors and specificity of ACX-362E

Limited treatment options and reports of reduced susceptibility to current treatment (11, 12, 45) emphasize the necessity for the development of novel antimicrobials. As CDI can be induced by use of broad spectrum antibiotics (7), new antimicrobials ideally should only target *C. difficile*, thereby maintaining integrity of the colonic microbiota. In this study, we have tested inhibitors HPUra and ACX-362E which specifically target the PolC enzyme, essential for DNA replication. The majority of PolC-inhibitors target gram-positive bacteria with low G+C content, but ACX-362E has been reported to demonstrate increased specificity towards *C. difficile* PolC *in vitro* and showed promising results for the efficacy *in vivo* based on a limited set of *C. difficile* strains (19, 20). The compound will progress to clinical trials in the near future (Acurx Pharmaceuticals, personal communication). The present study, is the largest survey of the efficacy of HPUra and ACX-362E against a large collection of clinical isolates consisting of many relevant PCR ribotypes to date. We have established that ACX-362E demonstrated lower inhibitory concentrations than the general gram-positive PolC-inhibitor HPUra in agar dilution experiments. The MIC_50_ and MIC_90_ of ACX-362E are similar to those of antimicrobials commonly used to treat *C. difficile* infection (metronidazole: MIC_50_ = 2 ug/mL and MIC_90_ = 4 ug/mL; vancomycin MIC_50_ = 2 ug/mL and MIC_90_ = 4 ug/mL (20); fidaxomicin = MIC_50_ = 0.125 ug/mL and MIC_90_ = 0.5 ug/mL (46)). We did not detect a significant difference in MICs between clades and ribotypes, demonstrating that PolC-inhibitors have the potential to be used as treatment for the majority - if not all - circulating *C. difficile* strains. This includes the epidemic types of PCR ribotype 027 and 078 (8, 47). These results are in line with other work that demonstrated only 2- to 4-fold differences in antimicrobial susceptibility between different clades for metronidazole, fidaxomicin and semi-synthetic thiopeptide antibiotic LFF571 (28). In the course of our experiments, we did not find any strains tha grew at the highest concentrations of either HPUra or ACX-362E tested. We cannot exclude rare pre-existing/intrinsic resistance, or the development of resistance in strains over time, for instance through multi-drug exporters, but at present there is no evidence for this. PolC-inhibitors are competitive inhibitors of polymerase activity by binding in the active site. Mutations that abolish binding of HPUra or ACX-362E are likely to affect the essential enzymatic activity of the polymerase and for that reason unlikely to occur *in vivo*. A single mutation (*azp-12*) has been described in *B. subtilis* that confers resistance to HPUra (48). This T>G transversion results in the replacement of a serine with an alanine in the highly conserved PFAM07733 domain of the polymerase (49). To our knowledge, it is unknown whether this mutation prevents binding of HPUra to PolC of *B. subtilis*. Few other mutations have been described that confer resistance against other PolC-inhibitors (50, 51). It will be of interest to see if similar mutations in *C. difficile* result in resistance to HPUra and/or ACX-362E and what the effect on binding of these compounds to *C. difficile* PolC is.

ACX362 may have off-target effects unrelated to its replication-inhibitory activity For *C. difficile*, it has not been established that PolC inhibition is the sole mode of action of the inhibitors. In our experiments, we found that *S. aureus* was sensitive to both HPUra and ACX-362E and even more so than *C. difficile*. It should be established if this is due to inhibition of PolC or mediated by an alternative mechanism. If ACX-362E targets DNA replication in *S. aureus*, we would also expect to find an increase in *oriC*:*terC* ratio upon ACX-362E exposure in this organism.ACX-362EACX-362E Alternatively, ACX-362E may also affect the activity of the other PolIII-type polymerase, DnaE, in *S. aureus*. PolIII-inhibitors can affect PolC, DnaE or both (51), though *in vivo* activity appears to correlate with PolC-inhibition. Both *C. difficile* and *S. aureus* possess PolC and DnaE polymerase, but the DnaE enzymes are of different families (DnaE1 in *C. difficile* and DnaE3 in *S. aureus*) (52). To verify the mode of action, and whether a ACX-362Edifferent DnaE-type polymerases explain the increased sensitivity of *S. aureus* compared to *C*. the activity of ACX-362E towards purified DnaE and PolC from both organisms should be determined.

Though it is clear that ACX-362E inhibits *C. difficile* efficiently and shows limited activity towards certain other anaerobes (19), these findings highlight the necessity to perform additional (microbiome) studies to more clearly define the antimicrobial spectrum of this compound. It also shows that ACX-362E may have therapeutic potential outside treatment of CDI.

### Regulators of the transcriptional response to PolC-inhibitors

The present study is the first to describe the transcriptional response of *C. difficile* to inhibition of DNA replication. We find that ~200 genes show differential expression under conditions of PolC-inhibition by both HPUra and ACX-362E, when compared to non-treated cells (**Supplemental Table 2, Figure 3**). When considering only ACX-362E, approximately 13% of all genes in the genome show statistically significant altered transcription. We demonstrate that this large reprogramming of transcription is likely to be caused directly by a gene dosage shift (**Figure 4 and 5**).

In addition to direct effects, it is conceivable that at least part of the transcriptional response is indirect. Our list of differentially regulated genes includes several putative regulators; sigma factors (including *sigE*, *sigG* and *sigH*), transcription factors and anti-terminators. The relatively long time (5h) at sub-MIC levels of antimicrobials may have contributed to secondary effects, through one or more of these regulators. Though shorter induction times are thought to provoke more compound-specific responses (53), we did observed a highly consistent transcriptional signature with both HPura and ACX-362E (**Supplemental Table 2**, **Figure 3**).

Major stress response pathways are poorly characterized in *C. difficile*. On the basis of experiments in other organisms (21-23, 54-56), we expect that inhibition of DNA replication inhibition might possibly induce an SOS response (LexA)(57), a DnaA-dependent transcriptional response (21), and possibly a heat shock response (HrcA/CtsR)(58) and/or a general stress response (σ^B^)(42). Of these, the best characterized stress response pathway in *C. difficile* is the one governed by σ^B^ (41). We noted a significant overlap in putatively σ^B^-dependent genes and those differentially expressed upon exposure to PolC-inhibitors (**Supplemental Table 2**). In addition, our luciferase reporter fusions directly implicate *sigB* in the expression and/or up-regulation of some of these (**Figure 7**). It should be noted that the *sigB-*operon itself was not differentially expressed in our RNA-seq analysis. A *sigB*-reporter fusion suggests that *sigB* is transiently up-regulated prior to the time point of sampling for the RNA-Seq analysis (**Figure 6**). Similar transient up-regulation of *sigB* followed by a persistent response of σ^B^-dependent gene expression has been observed in other organisms (58-60). To our knowledge, this is the first indication that *sigB*- and SigB-dependent gene expression could be subject to a gene-dosage effect.

To date, no genes have been identified that are regulated by DnaA in *C. difficile* and direct regulation of genes through the other stress response pathways has not been demonstrated. Putative LexA-regulated genes of *C. difficile* were identified *in silico* (61) and some of these (such as the *uvr* excinuclease and 30S ribosomal protein S3) were differentially expressed in our dataset (**Supplemental Table 2**). Indeed, we find that only part of the transcriptional up-regulation of *uvrB* as a result of ACX-362E exposure could be explained by *sigB* (**Figure 7D**), and LexA-dependent regulation might play a role. Pleiotropic phenotypes have been described for a *C. difficile lexA* mutant (62) and it is likely that other LexA targets than those identified *in silico* exist. No mutants of *hrcA* or *ctsR* have been described for *C. difficile*, but transcriptome and proteome analyses have been performed with heat shocked cells (42°C (60), or 41°C (59, 63, 64)). Similarities between these datasets and ours include genes encoding proline racemase (*prdF*), chaperones (*groEL*, *groES*), thioredoxin systems (*trxA*, *trxB*) and Clp-proteases (*clpC*, *clpP*).

Many parameters (such as the medium used, cell density, concentration of antibiotics, and protocol used to arrest transcription between cell harvest and lysis) can influence overall transcription signatures, and can also govern an incomplete overlap between our data and the stress regulons determined by others (53).

### Genome location contributes to the transcriptional response to PolC-inhibition

Our analysis of differential regulation in relation to genome location revealed a striking pattern of relative up-regulation for *oriC*-proximal genes, and down-regulation for *terC*-proximal genes under conditions of PolC inhibition (**Figure 4**). Antimicrobials directed at DNA replication in bacteria have a profound negative effect on the processivity of replication forks, though initiation of DNA replication is not or only marginally affected (23, 39). As a consequence, the presence of multiple replication forks simultaneously increases the copy numbers of genes located in close proximity of the origin of replication and such a gene dosage differences can result in functionally relevant transcriptional differences, either directly or indirectly (15). We found an increase of *oriC:terC* ratio when performing MFA on chromosomal DNA of cells subjected to a sub-inhibitory concentration of ACX-362E (and HPUra, albeit less pronounced) (**Figure 5**), consistent with findings in other organisms (23). This is the first demonstration of gene-dosage dependent transcriptional regulation in *C. difficile*.

An example of direct regulation by gene dosage can be found in *Vibrio*, for instance, where the location of ribosomal protein clusters close to the origin is crucial for fast growth, because increased copy number under condition of multi-fork replication allows for higher expression levels (65). We note that ribosomal gene clusters are up-regulated when DNA replication is inhibited in our experiments (**Table 2**, **Table 3**, **Supplemental Table 2**), suggesting that a similar mechanism may be active in *C. difficile*.

An example of indirect regulation as a result of gene dosage is the induction of competence genes in *S. pneumoniae* (23). Competence is believed to be a stress response in this organism, that lacks a canonical (σ^B^-dependent) stress response pathway. Key regulatory genes for competence development are located close to the origin, and replication inhibition therefore leads to the induction of origin distal competence genes (23). In our experiments, the large overlap with the proposed σ^B^ regulon (41), the origin proximal location of the *sigB* operon (8.5kb-10kb) (40) and the σ^B^-dependence of selected differentially expressed genes suggests that at least part of the transcriptional response to PolC-inhibition can be explained by an indirect gene dosage effect. The positioning of stress-response regulators close to *oriC* may therefore be a conserved strategy in bacteria to respond to DNA replication insults that is independent of the nature of the regulator.

Though it is likely that an increase in gene copy number leads to an increase in transcription of these genes, it is less clear whether this is the case for the observed down-regulation. Most methods of normalization for transcriptome analyses are based on the assumption that there is no overall change in transcription or that the number of transcripts per cells is the same for all conditions and this may not be the case when a global copy number shift occurs (15). Absolute transcript levels for down-regulated genes might therefore be similar under both conditions (but lower relative to *oriC*-proximal transcripts).

It is interesting that certain processes are highly enriched in the list of genes up-regulated under conditions of PolC-inhibition (most notably ribosome function and DNA-related functions), whereas this is less so for the down-regulated genes. This suggests that pathways susceptible to replication dependent gene-dosage effects demonstrate a functional clustering of genes near *oriC*, whereas clustering of genes from specific pathways in the *terC*-proximal region is less pronounced. Indeed, most ribosomal gene clusters in *C. difficile* are located close to the origin of replication (31, 40) and also many genes involved in DNA replication and repair are located in these regions. Consistent with this, positioning of genes involved in transcription and translation close to the origin appears to be under strong selection as such genomes tend to be more stable (66).

In conclusion, both direct and indirect effects of gene dosage shifts are likely to contribute to the transcriptional response of *C. difficile* to replication inhibition.

## Materials and Methods

### Bacterial strains and culture conditions

Plasmids and bacterial strains used in this study can be found in **Table 4**. Note that this table only contains laboratory strains; the clinical isolates used for the agar dilution experiments (see below) are listed in **Supplemental Table 1**. *E. coli* was cultured aerobically at 37°C (shaking at 200 rpm) in LuriaBertani (LB) broth supplemented with 20 µg/ml chloramphenicol and 50 µg/ml kanamycin when appropriate. *C. difficile* strains were cultured in Brain Heart Infusion (BHI) broth (Oxoid) supplemented with 0,5% yeast extract (Sigma-Aldrich), *Clostridium difficile* Selective Supplement (CDSS, Oxoid) and 20µg/ml thiamphenicol when appropriate. *C. difficile* was grown anaerobically in a Don Whitley VA-1000 workstation in an atmosphere of 10% CO_2_, 10% H_2_ and 80% N_2_. Liquid cultures were grown under gentle agitation (120 rpm).

**Table 4.** Plasmids and strains used in this study.

### Agar dilution

HPUra and ACX-362E were tested against a collection of *C. difficile* clinical isolates. 375 clinical isolates have been collected during the ECDIS study (6). All strains were characterized by PCR ribotyping (67) and by PCR to confirm the presence of genes encoding toxins A, B and binary toxin (68-70). Of the 375 clinical isolates, we excluded stocks that were found to contain more than one strain and isolates that could not be recultured. Testing was therefore performed on 363 isolates (**Supplemental Table 1**). *C. difficile* ATCC 700057, *B. fragilis* ATCC 25285 and *S. aureus* ATCC 29213 were used as controls (71).

The strains were tested for the different concentrations of antimicrobial using the agar dilution method according to Clinical & Laboratory Standards Institute guidelines (26, 27). In short, the antimicrobials were diluted into Brucella Blood Agar (BBA) supplemented with hemin and vitamin K1. Bacterial isolates were cultured on blood agar plates and after 24 hours re-suspended to a turbidity of 0.5 McFarland in phosphate buffered saline (PBS). The strains were inoculated onto BBA solid media containing the PolC-inhibitors using multipoint inoculators to a final concentration of 10^4^ CFU per spot. Each series of antimicrobial agents was tested from the lowest concentration to the highest concentration. Two control plates without antibiotics were inoculated to control for aerobic contamination and purity of anaerobic growth. At the end of the final series, two control plates were inoculated to verify the final organism viability and purity. Only experiments where both positive and negative controls performed according to expectations were included. Plates were incubated anaerobically in a Don Whitley VA-1000 workstation in an atmosphere of 10% CO_2_, 10% H_2_ and 80% N_2_ and the MICs were recorded after 24 and 48 hours and are presented in the manuscript with values at 48h, according to the CLSI guidelines (26).

### Sub-MIC determination

*C. difficile* 630Δ*erm* (30, 31) was grown in 20 mL Brain Heart Infusion (Oxoid) supplemented with 0.5% yeast extract (Sigma-Aldrich) (BHI/YE) starting from an optical density measured at 600nm (OD_600_) of 0.05 using an exponentially growing starter culture (3 biological replicates per concentration). To determine the effects on growth at sub-inhibitory concentrations of ACX-362E, cells were cultured in the presence of the following concentrations; 0.25, 0.5, 1, 2, 4, and 8 µg/mL ACX-362E and compared to an untreated culture. To determine the effects on growth at sub-inhibitory concentrations of HPUra, cells were cultured in the presence of the following concentrations: 10, 20, 40 µg/mL HPUra and compared to an untreated culture. The OD_600_ was monitored every hour until stationary phase was reached.

### Marker Frequency analysis

*C. difficile* 630Δ*erm* (30, 31) was grown in 20 mL BHI supplemented with 0.5% yeast extract with sub-MIC amounts of antimicrobials (HPUra: 35 μg/mL; ACX-362E: 4 μg/mL; metronidazole: 0.25 μg/mL; fidaxomicin: 0.00125 μg/mL, surotomycin: 0.625 μg/mL), starting from an OD_600_ of 0.05 using an exponentially growing starter culture. These samples were taken in the course of an independent, but simultaneously performed, set of experiments for which we obtained surotomycin and fidaxomicin from Cubist Pharmaceuticals. Metronidazole was commercially obtained (Sigma-Aldrich). We confirmed that these concentrations did not lead to a >30% reduction in growth compared to non-treated cultures (**Supplemental Figure 1** and data not shown). In parallel, cultures were grown without inhibitors from the same starter culture. All conditions were performed in biological triplicates. Previous experiments have shown that *ori*:*ter* differences are reliably detected >3h after dilution into fresh medium (23). Therefore, after 5 hours, 1mL cells was harvested (OD_600_ ~ 0.5), and stored at -20°C. Isolation of chromosomal DNA was performed the next day with the QIAamp DNA Blood Mini kit (Qiagen) according to the instructions of the manufacturer. Marker frequency analysis (MFA) was performed to assess the relative abundance of origin proximal genes relative to terminus proximal genes. As a proxy for *oriC*, a probe was designed that targets the CD0001 region (CD0001-probe-FAM). By using plots of the GC skew ([G - C]/[G + C]) generated using DNAPlotter (72), the approximate location of the terminus region for the *C. difficile* chromosome was determined and a probe targeting this region (CD1931) was designed (CD-1931-probe-TXR). Probe design was performed with Beacon Designer^TM^ (Premier Biosoft, Palo Alto CA, USA). Real-time PCR reactions were performed on a Biorad CFX96^TM^ real-time PCR detection system (95°C 15 m, 39 × (94°C 30 s + 55°C 30 s + 72°C 30 s). Sequences for primers and probes are listed in **Table 5**. For each biological replicate, three technical replicates were performed. Amplification efficiency was determined using standard curves obtained from DNA late stationary phase cells of strain 630Δ*erm*, for which an *oriC*:*terC* ratio of 1 was assumed. RT-PCR results from antibiotic treated cells were normalized to the *oriC*:*terC* ratio of DNA samples (3 biological replicates) from non-treated cells. Calculations were performed in Microsoft Office Excel 2010, plotted using Prism 7 (GraphPad) and prepared for publication in Corel Draw Suite X8. Significance was determined using a One-way ANOVA and a Tukey’s test for multiple comparisons (GraphPad).

**Table 5.** Oligonucleotides and probes used in this study.

### Growth and RNA isolation for RNA-Seq

For RNA-Seq analysis, *C. difficile* 630*Δerm* (30, 31) was grown for 5h in BHI medium with HPUra (35 ug/mL) or ACX-362E (4 µg/mL) starting from an OD_600_ of 0.05 using an exponentially growing starter culture, after which cells (3mL) were harvested for RNA isolation. These concentrations were some of the highest concentrations that resulted in a modest effects on growth (<30% reduction of growth compared to wild type cells, **Supplemental Figure 1**). RNA isolation was performed with NucleoSpin© RNA kit (Macherey-Nagel). Although the kit includes on column rDNAse digestion, a second treatment was performed in solution and RNA was precipitated and recovered by NaAc precipitation to remove residual DNA. Concentration determination and quality control (16S/23S ratio and RNA integrity number [RIN]) was performed with a fragment analyzer (Agilent bio-analyzer), according to the instructions of the manufacturer. Samples with a RIN>9 and 16S/23S ratio >1.4 were submitted for analysis by RNASeq.

### RNA-Seq

RNA-Seq was performed at a commercial provider (GenomeScan, Leiden, The Netherlands). In short, the NEBNext Ultra Directional RNA Library Prep Kit for lllumina was used to process the samples. Sample preparation was performed according to the protocol "NEBNext Ultra Directional RNA Library Prep Kit for lllumina" (NEB #E7420S/L). Briefly, after selective removal of rRNA (Ribo-Zero rRNA Removal Kit for gram-positive Bacteria) and fragmentation of the mRNA, cDNA synthesis was performed. cDNA was ligated to the sequencing adapters and the resulting product was PCR amplified. Clustering and DNA sequencing using the lllumina NextSeq 500 platform was performed according to manufacturer’s protocols. A concentration of 1.5 pM of DNA was used. Image analysis, base calling, and quality check was performed with the lllumina data analysis pipeline RTA vl.18.64 and Bcl2fastq v2.17. Per sample, four technical replicates were included in the RNA-Seq experiment. In case of insufficient reads, the sample was re-run on another flow cell to reach satisfactory quantities (≥ 20 M).

### Analysis of RNA-Seq data

Analysis of the data was performed using T-REx, a user-friendly webserver which has been optimized for the analysis of prokaryotic RNAseq-derived expression data (73). The pipeline requires raw RNA expression level data as an input for RNA-Seq data analysis. For data normalisation and determination of the genes, the factorial design statistical method of the RNA-Seq analysis R-package EdgeR is implemented in the T-Rex pipeline. Some samples displayed incomplete rRNA depletion and rRNA mapping reads had to be removed manually prior to analysis.

To analyse the genome-wide pattern in differential gene expression a sliding window analysis was performed essentially as described (23). In short, genome locations (start of the locus tag) were coupled to the locus tags in the T-Rex output. Next, the median log(FC) was calculated for bins of 51 loci with a stepsize of 1. For each bin of [X_1_, X_2_…X_51_] the median absolute deviation of the median (MAD=median(|X_i_-median(X)|) was calculated as an robust indication of the distribution around calculated median values. Calculations were performed and three curves (median, median-MAD and median+MAD) were plotted in Microsoft Office Excel 2010 and the graph was prepared for publication using Adobe Photoshop CC and Corel Draw Suite X8.

A GSEA analysis (33) was performed via the Genome2D webserver (73), using our reference genome sequence for *C. difficile* 630Δ*erm*, Genbank identifier LN614756.1 (listed in Genome2D as “Clostridioides_difficile_630Derm”)(31). As input a single list of locus tags was used of either up- or down regulated genes. The output was copied to Microsoft Excel 2010. The single_list column was split, and a column was inserted to calculate the significance of the overrepresentation using the formula “(# hits in list/ClassSize)*-log(p-value; 2)” to allow for sorting of the output of the GSEA analysis by significance.

Data for the RNA-Seq experiment has been deposited in the Gene Expression Omnibus (GEO) database (74), accession number GSE116503.

### General molecular biological techniques

*E. coli* strain DH5α was used for maintenance of all plasmids. All plasmid transformations into *E. coli* were performed using standard procedures (75). *E. coli* CA434 was used as a donor for conjugation of plasmids into the recipient *C. difficile* (76). Conjugation was performed as previously described (76). Briefly, 1 mL of an overnight culture of donor cells was mixed with 200 µl of the recipient and spotted onto anaerobic BHI agar plates and incubated for 5-8 hours. After incubation cells were collected and tenfold serial dilutions were plated onto fresh BHI plates containing thiamphenicol and CDSS.

Plasmid DNA was isolated using the Nucleospin Plasmid (Macherey-Nagel) mini prep kits per manufacturer’s instructions. *C. difficile* genomic DNA was isolated using the DNeasy Blood and Tissue kit (Qiagen) with pre-treatment for gram-positives according to instructions of the manufacturer.

### Construction of luciferase-reporter fusion plasmids

All PCR reactions for plasmid construction were carried out with Q5 polymerase (New England Biolabs). Putative promoter regions were amplified using *C. difficile* 630Δ*erm* chromosomal DNA (31) as a template.

The PCD3412 luciferase reporter plasmid was created by restriction-ligation using the restriction enzymes *KpnI* and *SacI*. P_CD3412_ was amplified using primers oIB-26 and oIB-27 (**Table 5**). The resulting dsDNA fragment was digested and ligated into KpnI-SacI digested pAP24 (44), yielding plasmid pIB27. Plamids pIB68 (P_CD0350_), pIB69 (P_CD2963_), pIB74 (P_CD3614_) and pPH28 (P_CD0007_) were constructed using a Gibson assembly (77). The plasmid backbone of pAP24 was linearized by PCR using primers oWKS-1580/oWKS-1582, and the predicted promoter areas of CD0350, CD2963 and CD3614 were amplified with primers oIB-80/oIB-94, oIB-82/oIB-95, oIB-92/oIB-100 and oPH-19/oPH-20, respectively. Primers were designed using the NEBuilder assembly tool v.1.12.17 (New England Biolabs) using a 30 bp overlap. For the assembly, 100 ng of vector DNA was assembled to a five-fold molar excess of the PCR fragment of the desired promoter using a homemade Gibson Assembly Master Mix at 50°C for 30 minutes (final concentration: 4 U/µl Taq Ligase (Westburg), 0.004 U/µl T5 exonuclease (New England Biolabs), 0.025 U/µl Phusion polymerase (Bioké), 5% polyethyleneglycol (PEG-8000),10 mM MgCl_2_, 100 mM Tris-Cl pH=7.5, 10 mM dithiothreitol, 0.2 mM dATP, 0.2 mM dTTP, 0.2 mM dCTP, 0.2 mM dGTP, and 1 mM β-nicotinamide adenine dinucleotide) and transformed into *E. coli* DH5α. Transformants were screened by colony PCR using primers oWKS-1240 and oWKS-1241. Transformants yielding PCR fragments of the correct size were verified by Sanger sequencing.

### *Construction of* C. difficile *IB56 (ΔsigB)*

The up- and downstream regions (both 950 bp each) of the *sigB* coding sequence were amplified with primers oIB-44/oIB-45, and oIB-46/oIB-47, respectively. Vector pMTL-SC7315 (43) was linearized by PCR using primers oWKS-1537/oWKS-1538. Assembly was done according to the method of Gibson (44, 77). The assembled plasmid was transformed into *E. coli* DH5α and verified using PCR and Sanger sequencing. Generation of the unmarked *sigB* deletion mutant was performed using Allele-Coupled Exchange essentially as described (43, 78). Briefly, pIB54 was introduced into *C. difficile* 630Δ*erm* (31) by conjugation. Transconjugants were grown for two days on BHI agar plates supplemented with yeast extract, thiamphenicol and CDSS, struck onto fresh pre-reduced plates and incubated anaerobically at 37°C for two days. Single-crossover integration was confirmed using PCR, and those clones were plated onto non-selective BHI agar plates to allow the second-crossover event to occur. Colonies were then serially diluted and plated onto minimal agar supplemented with 50 µg/ml 5-fluorocytosine (Sigma) as described (43). DNA was isolated from thiamphenicol-susceptible colonies and the chromosomal deletion was verified by PCR using primers oIB-78/oIB-79 (**Supplemental Figure 3**) as well as Sanger sequencing of the PCR product using primers oIB-53/oIB-76/oIB-78.

### Luciferase reporter assay

Strains containing luciferase reporter plasmids were inoculated to an OD_600_=0.05 from an exponentially growing starter culture. Fresh inoculums were grown in BHI broth supplemented with yeast extract, with or without 4 µg/ml ACX-362E for 5 (for putative *sigB* target genes) or 5.5 (*sigB* promoter) hours. The supernatants from 1mL of culture (harvested by centrifugation for 10 min, 4°C, 8000 rpm) were analyzed in a GloMax®-Multi Microplate Multimode Reader (Promega) as described before (44). Statistical significance of the data (p<0.05) was determined by two-way analysis of variance (ANOVA) and a pairwise Tukey-Kramer test using Prism 7 (GraphPad) where appropriate.

#### Accession numbers

GEO GSE116503; GenBank LN614756.1

## Author’s contributions

EVE, IMB, EJK and WKS designed experiments. GW and EJK provided reagents or strains. EVE, IMB and IMGJBS performed experiments. EVE, IMB and WKS analyzed data. EVE, IMB and WKS wrote the manuscript. All authors read and approved the manuscript.

## Acknowledgements

This work was supported, in part, by a VIDI fellowship (864.13.003) from the Netherlands Organization for Scientific Research, a Gisela Thier Fellowship from the Leiden University Medical Center and intramural funds to WKS. Anne de Jong is acknowledged for expert assistance with the RNA-Seq analysis, Els Wessels for assistance in designing the MFA real-time PCR and Jelle Slager for helpful suggestions for the sliding window analysis.

## Supplemental information

**Supplemental Table 1**. Characteristics and minimal inhibitory concentrations of the clinical isolates used in the agar dilution experiments.

**Supplemental Table 2**. Lists of the genes that are differentially expressed in the presence of PolC-inhibitors compared to non-treated cells. **A**. HPUra differentially expressed genes. **B**. ACX-362E differentially expressed genes.

**Supplemental Table 3**. Overlap with published σ^B^ regulon. **A**. σ^B^ -dependent stress response genes differentially regulated in the ACX-362E dataset. **B**. Genes containing a σ^B^ consensus promoter differentially regulated in the ACX-362E dataset.

**Supplemental Figure 1**. Inhibition of growth by varying concentrations of PolC-inhibitors.

**Supplemental Figure 2**. Chloramphenicol does not result in a gene dosage shift.

**Supplemental Figure 3.** Confirmation of the *sigB* mutant strain (IB56).

Supplemental information is also available from Figshare (10.6084/m9.figshare.7117463).

